# Performance of bitumen coating sheet using biomass pyrolysis oil

**DOI:** 10.1101/592246

**Authors:** Yanru Ren, Lei Zhang, Wenfeng Duan, Jia Guo, Zhongqiang Han, Michael D. Heydenrych, Aijun Zhang, Kaili Nie, Tianwei Tan, Luo Liu

**Author notes:** Author of Correspondence: Luo Liu.

## Abstract

The “green” production of bitumen has raised increasing interest in recent years to reduce the environmental, energy and petro-based concerns. Bio-oil, prepared by biomass pyrolysis, can be used as substitute for petro-based bitumen in bitumen or bitumen-based coatings, for its similar properties of good adhesion and anti-corrosion characteristics. Although biomass is a renewable and widespread chemicals resource, its value-added utilization is still difficult. Several studies have qualitatively demonstrated the use of bio-bitumen in practical applications. The present study investigates the effects and properties and the incorporation of bio-bitumen shown to improve the performance of traditional petro-bitumen to some extent. Bio-bitumen was prepared from biomass pyrolysis oil and applied to self-adhesive and doped hot-melt sheets. Resulting physical properties demonstrate that bio-bitumen is a potential substitute in bitumen coating sheet.

**IMPLICATIONS:** This paper is intended to verify the effect of pyrolyzed bio-oil from wheat straw on the performance of bitumen, as well as the feasibility of application in the coating sheet. Up to now, the research on bio-bitumen is mainly in pavement bitumen. In the present research, bio-bitumen was applied to the coating sheet in different proportions. Interestingly, the prepared coating sheet exhibited higher adhesion. Other performances, such as temperature stability, mechanical strength and temperature flexibility of coating sheet showed improvement in the presence of bio-oil, which indicated the suitability of bio-oil in coating sheet bitumen.

## INTRODUCTION

As the most widely used coating and pavement cementing material, bitumen is in great annual demand. In recent years, with the increasing shortage of crude oil resources, with enhanced environmental concerns, and with a shift in refinery strategies to a reduced pitch (bitumen) production, the bitumen industry is facing unprecedented challenges and the need to reduce the application of petroleum bitumen in the construction industry is a hot industrial and research topic. Biomass-based alternatives could be a valid and valuable development. Biomass refers to the various organisms formed by photosynthesis, including animal, plant and microorganism biomass. As the fourth largest energy resource, biomass is abundant and the most important raw material for producing bio-oil. At present, researchers have extracted bio-oil from microalgae (Brennan et al. **2010**), manure (Fini et al. **2011**), oil seeds (Kong et al. **2019**; Kong et al. **2018**; Kong et al. **2017**), short rotation forestry wood (Prayogo et al. **2014**; Van et al. **2010**), corn stover (Raouf et al. **2010**), rice husks (Zheng et al.**2010**), and other materials (Van et al. **2010**; Sanna et al. **2011**). Many innovative techniques for producing bio-oil from biomass have been developed, mostly including thermochemical techniques such as hydrolysis, pyrolysis and liquefaction (Fernandez et al. **2019**; Van et al. **2008**; Van et al. **2006**; Demirbas **2001**; Önal et al. **2011**), with pyrolysis as the most widely used technology. In the absence of air, pyrolysis converts organic matter into a solid biochar, a condensable bio-oil and non-condensable small molecule gases (Goyal et al. **2008**; Akhtar et al. **2010**). Pyrolysis can occur in different regimes according to the residence time of the solid and gas phases in the reactor, with fast, intermediate and slow pyrolysis having been investigated (Van et al. **2010**; Demirbas et al. **2001**). Fast pyrolysis is widely used due to its simplicity, high bio-oil efficiency, and low cost (Van et al. **2010**; Van et al. **2018**).

A cementitious material obtained by incorporating a given fraction of a pyrolytic bio-oil into a conventional petroleum pitch is called a bio-bitumen. Several researchers have studied the practical application of bio-bitumen (Bridgwater et al. **2000**). Adding bio-oil to matrix bitumen can change its temperature sensitivity (Önal et al. **2011**) and modify the overall properties (Yang et al. **2014**; Zargar et al. **2012**). Forest waste bio-bitumen can improve the high temperature performance of petro-based bitumen, although it has an adverse effect on the low temperature performance of the adhesive (Yang et al. **2013**). On the contrary, bio-bitumen produced from pig manure and edible oil has an inferior high temperature performance than traditional bitumen, but can improve low temperature performance (Fini et al. **2011**, Yang et al. **2017**; Wen et al. **2013**; Su et al. **2018**). The use of bio-bitumen not only reduces production costs, but more importantly it has a positive impact on the environment and can improve the performance of traditional bitumen to some extent.

Bitumen is also widely used in coating sheet due to its good adhesion, plasticity, water resistance, corrosion resistance and durability. Coating sheets based on bitumen are widely used in tunneling, roofing of industrial and private buildings, or as anticorrosive protection of metal pipes. In the present research, bio-bitumen is applied at different fractions into two types of bitumen sheets. Results on temperature stability, mechanical strength and temperature flexibility will demonstrate that bio-oil is suitable for the preparation of coating sheet bitumen.

## MATERIALS AND METHODS

### Materials

Two petro-based bitumen binders with 70 and 200 penetration grades (as determined by standard needle test at 25 °C, and denoted as 70# and 200# grades) were used in our study and were obtained from Sinopec (Jiangsu, China). The SBS (Styrene-Butadiene-Styrene) modifiers used in this study are 6302H or 4402 linear structures produced by Lichangrong chemical industry company (Hunan, China). The SBR (Styrene-Butadiene-Rubber) modifier is produced by Lichangrong chemical industry company (Hunan, China). The C5 resin used is produced by Exxon Mobil (Singapore).

### Preparation of bio-oil

The pulverized wheat straw (< 1 mm) was placed in a N_2_-purged tubular reactor and reacted at 500 °C for 15 s. The obtained cracking products are solid biochar, liquid bio-oil, and non-condensable gases. When using wheat straw as biomass species at the specified temperature and reaction time, the yield of bio-oil is 43%. Biochar represents 25% and the balance consists of non-condensable gases.

The pyrolysis oil was analyzed by Fourier Transform Infrared Spectrometer (Vertex 70V, Bruker). Since the prepared bio-oil was viscous, it was dissolved in dichloromethane followed by dropping the solution onto a KBr table. The sample was scanned within a test spectrum range of 400 to 4000 cm^−1^ after evaporating the solvent.

### Preparation of bio-bitumen coating sheet

The wheat straw pyrolysis oil (water content 8%) was applied directly to substitute part of the petro-bitumen into self-adhesive coating sheets and Styrene-Butadiene-Styrene (SBS) hot-melt coating sheets. No any pretreatment was carried out before use, except in some case it was distillated at 100 °C for dewatering.

### Bio-bitumen applied in self-adhesive coating

Self-adhesive coating sheets were prepared by firstly adding the petro-bitumen binders 70# and 200# into a mixer with mechanical stirring. After heating and melting at 120 °C for 1h, 5 wt% of Styrene-Butadiene Rubber (SBR) was added to the mixture. The temperature increased to 180 °C in a controlled manner (1 °C/min). SBS was then introduced into the mixture, with stirring for 2 h at 180 °C. Finally, bio-oil was added into the molten mixture at 5, 10, 15 and 20 wt%, and stirred for an additional 2 hours. The molten high temperature sample was quickly discharged and further used for testing after cooling.

### Bio-bitumen applied in SBS hot-melt coating sheet

SBS hot-melt coating sheets were prepared by firstly mixing the bitumen binders 70# and 200# under heating till 150 °C for 1 h. 10 wt% SBS were added and stirred at 180 °C for 3 hours. Again, bio-oil was added at 5, 10, 15 and 20 wt% while stirring for another 2 hours. The molten high temperature sample was quickly discharged and further used for testing after cooling.

### Performance tests of bio-bitumen modified coating sheets

#### Softening point

The softening point of the coating was determined by a bitumen softening point analyzer (WSY-025C, accuracy ±0.5°C), according to GB/T 4507-2014 (“Bitumen softening point determination method”), at a heating rate of 5 °C/ min.

#### Low temperature flexibility

According to GB/T 328.14-2007 test method for building coating sheets, the low-temperature flexibility of the said sheets is used to determine the low-temperature properties and resistance of bitumen coating sheets. The specimen size is (150 ± 1) mm × (25 ± 1) mm, and the temperature is maintained for 1 h ± 5 min in a low-temperature refrigerator (Aucma, accuracy ± 2 °C) in temperature range -20°C to -40 °C, and then tested with a low-temperature flexible tester (DWR-2). The temperature without visible cracks on the sample was recorded.

#### Peeling strength

The peeling strength was determined by Microcomputer-controlled electronic universal testing machine (MTS, CMT 6104), in accordance with GB/T 328.20-2007 (“Building waterproofing coating sheet test method Part 20: Bitumen coating sheet joint peeling performance where the stretching rate is 100 ± 10 mm/min at ambient temperature of 23 ± 2 ° C”). Peeling strengths were determined for application of the bio-bitumen coating sheets on both a petro-based coating sheet and on an aluminum plate, respectively.

#### Viscosity

The viscosity of bio-bitumen at 150 °C and 180 °C was measured by Rotating viscometer (Brookfield), using 26 rotors.

#### Heat resistance

The heat resistance was measured by Electric thermostatic drying oven (FXB-2, accuracy ± 2 ° C), according to GB-T 328.11-2007 (“Building coating sheet test method Part 11: Bitumen waterproofing membrane heat resistance to characterize coating materials heat resistance”). The heat resistances of the bio-bitumen at 70 °C (coating material for self-adhesive sheets) and 105 °C (coating material for hot-melt sheets) were characterized.

#### Density

The density was measured by Density balance (OHAUS, precision 1 mg). Triplicate measurements were performed, and averages are further reported (with deviations of triplicate samples within 10 kg/m^3^ from the averages).

#### Hardness

A Shore A hardness analyzer (TB51-LX-A) was used, according to GB/T 531.1-2008 (“Vulcanized rubber or thermoplastic rubber indented hardness test method Part 1 Shore hardness test method to test modified bitumen hardness”). The median value was measured five times on the surface of the sample, and average median results are further reported.

#### Maintained stickiness

Under the condition of (23±2 °C), the test piece was adhered to two clean and smooth mirror stainless steel plates, and repeatedly pressed three times with a pressure bar, according to GB/T 23441-2009. After 24 hours, it was suspended vertically and a 1 kg weight was suspended at the lower end of the lower plate, and the time (min) at which the test piece was completely peeled off from the upper plate was recorded.

## RESULTS AND DISCUSSION

### FTIR results

A sample of the bio-oil is illustrated in Figure 1. It is a dark and sticky liquid. It was analyzed by FTIR analysis, and the obtained infrared spectrum is shown in Figure 2.

**Figure 1.**
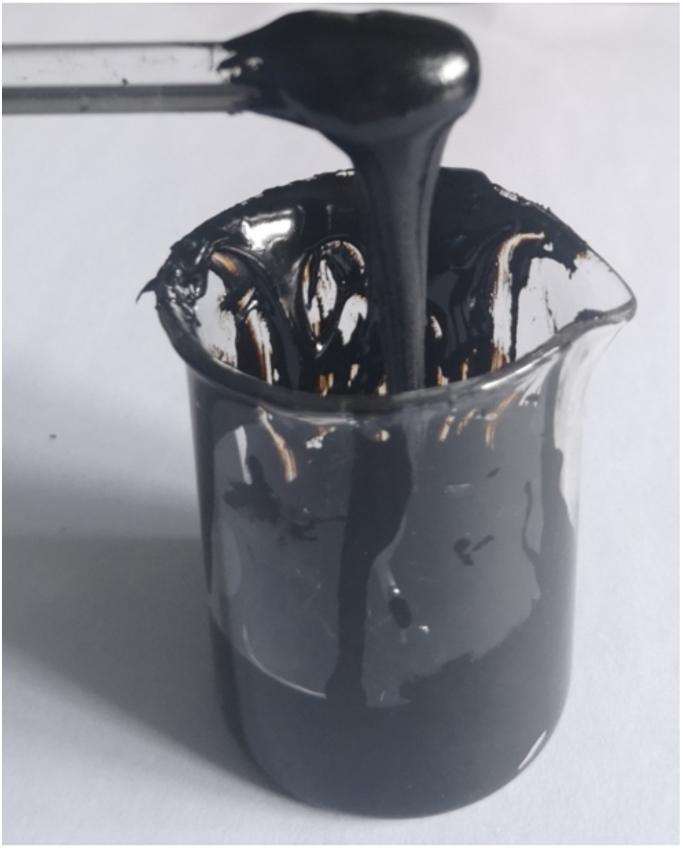
Sample of bio-oil obtained from pyrolysis of wheat straw.

**Figure 2.**
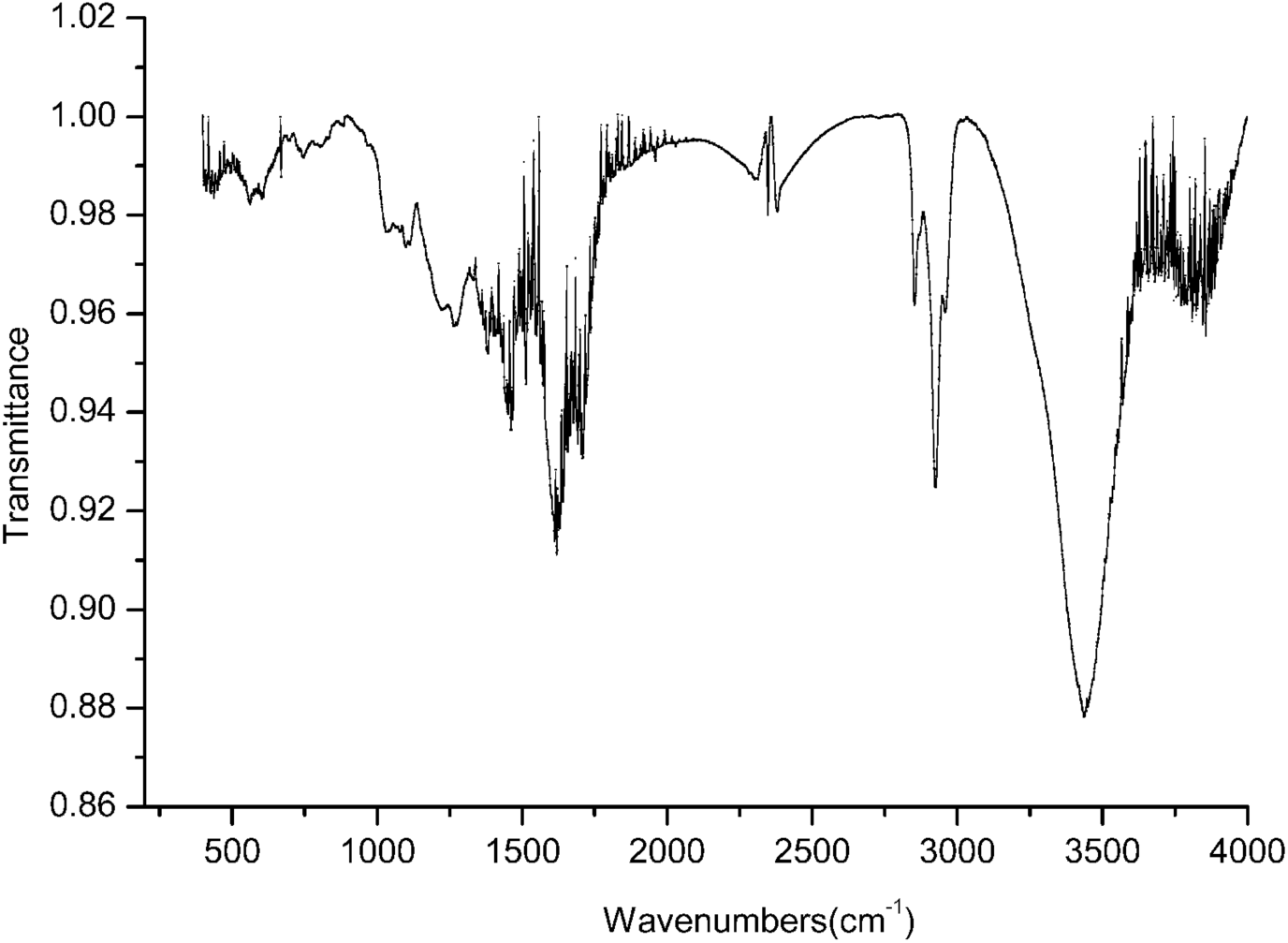
FTIR spectra of bio-oil from pyrolysis.

The bio-oil produced by pyrolysis of biomass is very complex, mainly containing water, acids, aldehydes, ketones, esters, furans, sugars and phenols [24]. The FTIR spectrum of Fig. 2 shows the similar result. It can be seen from Fig. 2 that the molecular structure of the sample is mainly composed of chain hydrocarbon derivatives; The C=C bond vibration characteristic peak near 1600cm^−1^ is more obvious, indicating that the sample contains more unsaturated bonds. A characteristic peak around 1700 cm^−1^ indicates that the sample contains a part of the carbonyl structure (C=O stretching vibration). The weak signal peaks around 1200 cm^−1^ and 3000 cm^−1^ indicate low aromatics content. Moreover, the absorption peak around 3450 cm^−1^ indicates the water contained in the sample due to undying during sample preparation.

### Bio-oil as substitute of petro-based bitumen

#### Effect of bio-oil directly added to a bitumen coating sheet

It can be seen from both table 1 and table 2 that the softening point is obviously increased with the increase of bio-oil addition, and the peeling strength of also increased for both the self-adhesive or SBS hot-melt coating sheets. In particular, the peel strength of the self-adhesive web was increased by 68% compared to the blank test when the amount of bio-oil added was 15.7%. The peel strength of SBS hot-melt coating sheets decreased when the amount of bio-oil added was 5%, but increased when the amount exceeded 10%. Moreover, the addition of bio-oil also slightly improved the low temperature flexibility of the self-adhesive coating sheet, but it adversely affected the low temperature flexibility of the SBS hot melt sheet, and the viscosity of the SBS hot melt bitumen coating sheet significantly increased.

**Table 1.** Effect of bio-oil directly added to self-adhesive coating sheets (using grade 70# matrix).

| Test items | Bio-oil wt% added |  |  |  |  |  |
| --- | --- | --- | --- | --- | --- | --- |
|  | 0% | 6.2% | 6.5% | 13.1% | 15.7% | 20% |
| Softening Point, °C | 80 | 82 | 87 | 86 | 91 | 90 |
| Heat resistance <sup>a</sup>, °C | 70 | 70 | 70 | 70 | 70 | 70 |
| Low temperature flexibility <sup>b</sup>, °C | -31.8 | -31.8 | -35 | -35 | -29 | -29 |
| Viscosity, cP | 4471 | 5463 | 4153 | 4750 | -- | -- |
| Density, kg/m<sup>3</sup> | 1150 | 1230 | 1160 | 1160 | 1150 | 1150 |
| Peel strength, N/mm | 1.83 | 2.29 | 2.18 | 2.29 | 3.08 | 2.95 |

<sup>a</sup>: heat resistance was measured at 70 °C and samples were not free-flowing; <sup>b</sup>: values indicate the temperature without visible cracks

**Table 2.** Effect of bio-oil directly added to SBS hot-melt sheets

| Test items | Bio-oil wt% added |  |  |  |  |  |
| --- | --- | --- | --- | --- | --- | --- |
|  | 0% | 5.5% | 6.0% | 11.4% | 11.2% | 11.5% |
| Softening Point, °C | 94 | 86 | 94 | 96 | 103 | 101 |
| Heat resistance <sup>a</sup>, °C | 105 | 105 | 105 | 105 | 105 | 105 |
| Low temperature flexibility <sup>b</sup>, °C | -34.9 | -28 | -29 | -28.3 | -28 | -29 |
| Viscosity, cP | 5339 | 7325 | 12380 | 10640 | 14700 | 10500 |
| Density, kg/m<sup>3</sup> | 1220 | 1230 | 1210 | 1230 | 1230 | 1220 |
| Shore A hardness | 3.52 | 4.02 | 4.48 | 5.24 | 4.24 | 4.0 |
| Peel strength, N/mm | 2.37 | 1.19 | 2.48 | 2.45 | 2.67 | 2.42 |

<sup>a</sup>: heat resistance was measured at 105 °C and no significant flow (< 2 mm) of samples was observed; <sup>b</sup>: values indicate the temperature without visible cracks

Bio-oil has a higher oxygen content than petroleum, so the aging rate is fast, resulting in an increase in the hardness of the bio-bitumen. This may also be the main reason for the increase in softening point. In addition, bio-oil also produces a small amount of coke during the preparation and aging process that can act as an aggregate, which is beneficial to the improvement of softening point and peel strength. It has also been observed that the incorporation of bio-oil significantly enhances the low temperature flexibility of self-adhesive coating sheets, as explained by the fact that bio-oil contains many C=C double bonds. Since the biochar particles have a high thermal stability, their presence can also help to alleviate the thermal stress caused by a low temperature.

#### Effect of bio-oil addition to a bitumen coating sheet after dewatering

Bio-oil obtained from biomass pyrolysis contain a considerable amount of moisture produced from dehydration reactions or that obtained from the moisture inherent in the biomass itself [7,8,25]. It is hence necessary to study the effect of moisture on modified bitumen coating sheets.

Table 3 shows the performance of adding bio-oil to the self-adhesive coating sheet after dewatering. When the addition amount is within 15%, the low-temperature flexibility of both mixes and of the blank test do not significantly differ. When the addition amount is 5% and 10%, the peel strength of the sheet-to-sheet and the sheet-to-aluminum plate is improved, but when the addition amount exceeds 10%, the softening point thereof is largely lowered. In addition, the Maintained stickiness of the coating sheet can be increased to about 2 hours as the amount of added bio-bitumen increases.

**Table 3.** Effect of bio-oil after water removal added to self-adhesive sheets of #70 grade

| Test items |  | Bio-oil wt% added |  |  |  |  |  |
| --- | --- | --- | --- | --- | --- | --- | --- |
|  |  | 0% | 5% | 10% | 15% | 20% | 30% |
| Soluble content, % |  | 75% | 75% | 78% | -- | -- | -- |
| Softening Point, °C |  | 99 | 98 | 90 | 92 | 88 | 84 |
| Heat resistance <sup>a</sup>, °C |  | 70 | 70 | 70 | 70 | 70 | 70 |
| Low temperature flexibility <sup>b</sup>, °C |  | -36 | -34.1 | -35 | > -35 | -30.5 | < -30 |
| Viscosity, cP | 150 °C | 30100 | -- | 23310 | 30300 | -- | -- |
|  | 180 °C | 9900 | 10540 | 11230 | 10070 | 11190 | 10570 |
| Density, kg/m<sup>3</sup> |  | 1110 | 1120 | 1150 | 1140 | 1140 | 1166 |
| Shore A hardness |  | -- | -- | 4.92 | 5.08 | 6.16 | 5.94 |
| Peel strength (N/mm) | Sheet-on-sheet | 2.23 | 2.25 | 2.88 | 2.33 | 2.64 | 2.40 |
|  | Sheet-on-aluminum | 1.54 | -- | 2.12 | 1.77 | 1.68 | 1.67 |
| Maintained stickiness (min) |  | 97 | 101 | 118 | -- | -- | -- |
| Self-adhesive bitumen re-peel strength (N/mm) |  | 1.29 | -- | 1.81 | -- | -- | -- |

<sup>a</sup>: Pure biomass oil soluble content after removal of water is 79%; <sup>b</sup>: Temperature values without observed cracks.

Table 4 shows that after dewatering, the softening point, peel strength and low temperature flexibility show a decreasing trend with the increase of the added amount. And under the same amount of addition, the softening point of the bio-bitumen is lower than that before dewatering. However, the peel strength is higher than that before dewatering. In addition, the incorporation of bio-oil also increases the viscosity and hardness of the bitumen. And after dewatering, the hardness is significantly greater than that before dewatering, the viscosity is significantly lower than that before dewatering. Overall, self-adhesive sheets have the best performance when bio-oil is without dewatering and added with 10%.

**Table 4.**
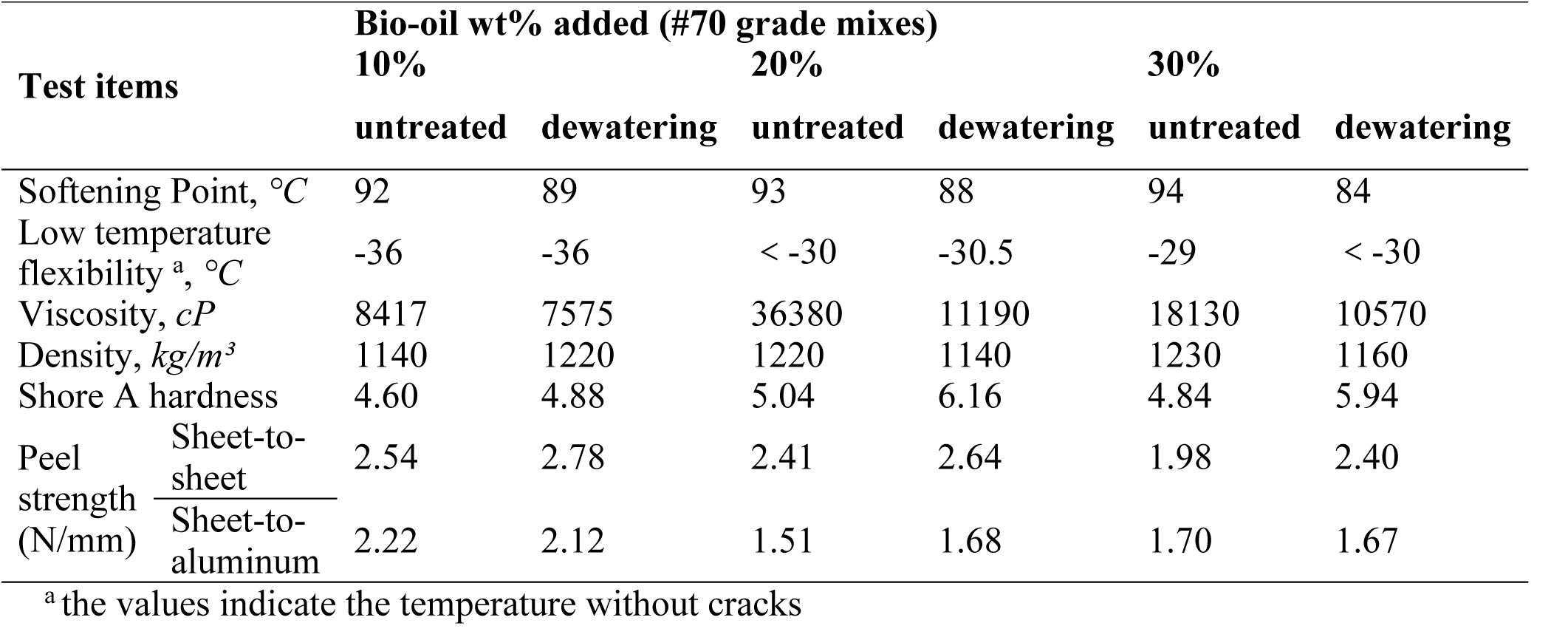
Effect of bio-oil added to self-adhesive coating sheets before and after dewatering

Table 5 shows the performance of added bio-oil to SBS hot-melt coating sheets after heating and dewatering. As the amount of added bio-oil increases, the low-temperature flexibility decreases. The peel strength decreased at 5% addition, and increased again at 10%, but remained below the blank value. In general, the bio-oil dewatering has an adverse effect on the performance of both coating sheets. This phenomenon may be due the fact that removing water destroyed the hydrogen bonds that help maintain structural stability.

**Table 5.**
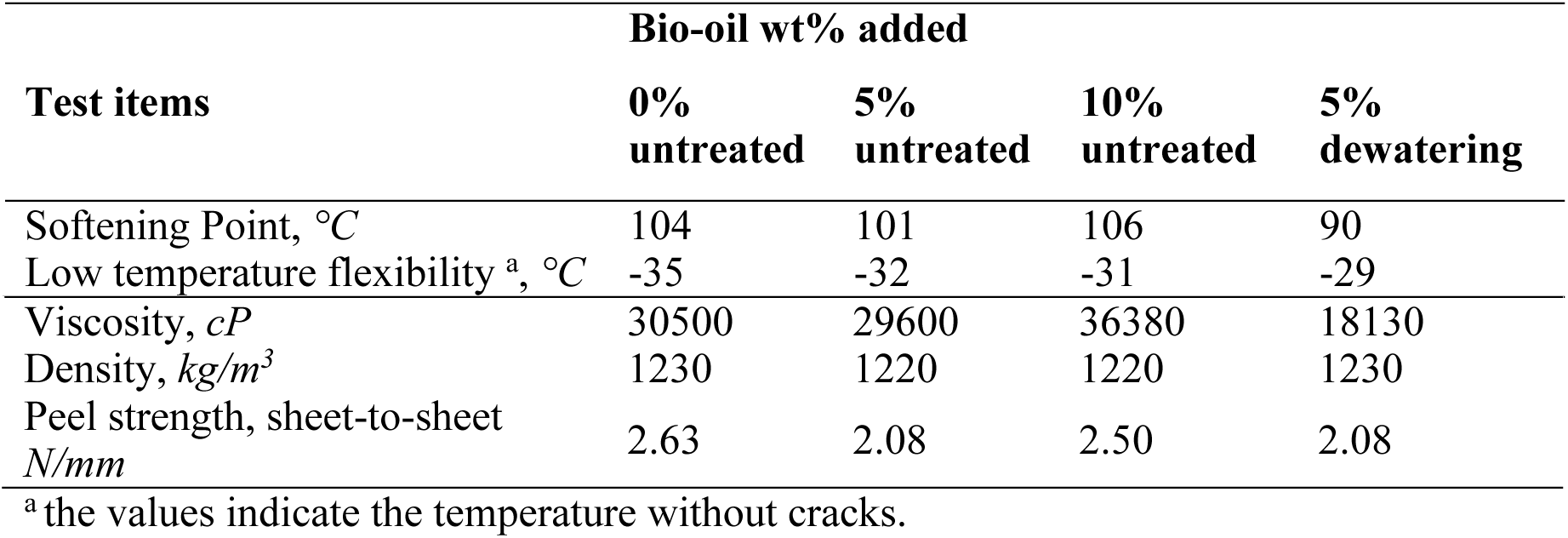
Effect of bio-oil added to SBS hot-melt sheets after dewatering

### Performance of coating sheet after aging

Table 6 shows the performance of the self-adhesive coating sheet after aging. Before heat aging test, the coating sheet with 10% bio-bitumen shows significantly higher peel strength than the blank. The peel strength decreased after UV aging more than twice its value after heat aging. After the heat aging test and the UV aging experiment, the peel strength and low temperature flexibility of the blank test and bio-bitumen are reduced to the same level.

**Table 6.** Performance changes after self-adhesive coating sheet aging test

| Test items |  | Bio-oil wt% added |  |
| --- | --- | --- | --- |
|  |  | 0% | 10% |
| Sheet-to-aluminum sheet<br>peel strength,<br>*N/mm* | Before heat aging | 1.54 | 2.12 |
|  | After heat aging <sup>a</sup> | 0.69 | 0.68 |
|  | After UV aging <sup>a</sup> | 0.34 | 0.36 |
| Low temperature<br>flexibility <sup>b</sup>, | Before heat aging | -36 | -36 |
|  | After heat aging | -32 | -33 |

<sup>1</sup> the heat aging condition is 168 h at 70 °C, and the ultraviolet aging condition is 45 °C for 240 h;
<sup>2</sup> the values indicate the temperature without cracks

#### Bio-oil replaces part of 70# and 200# bitumen in SBS hot-melt sheets

Table 7 shows that when partially replacing 70# bitumen, the low temperature flexibility is improved with the increase of the substitution ratio, however, not for the substitution of 200# bitumen. Regardless of the replacement of 70# or 200#, the softening point is significantly reduced when the substitution ratio is 20% compared to the 10% replacement rate.

**Table 7.**
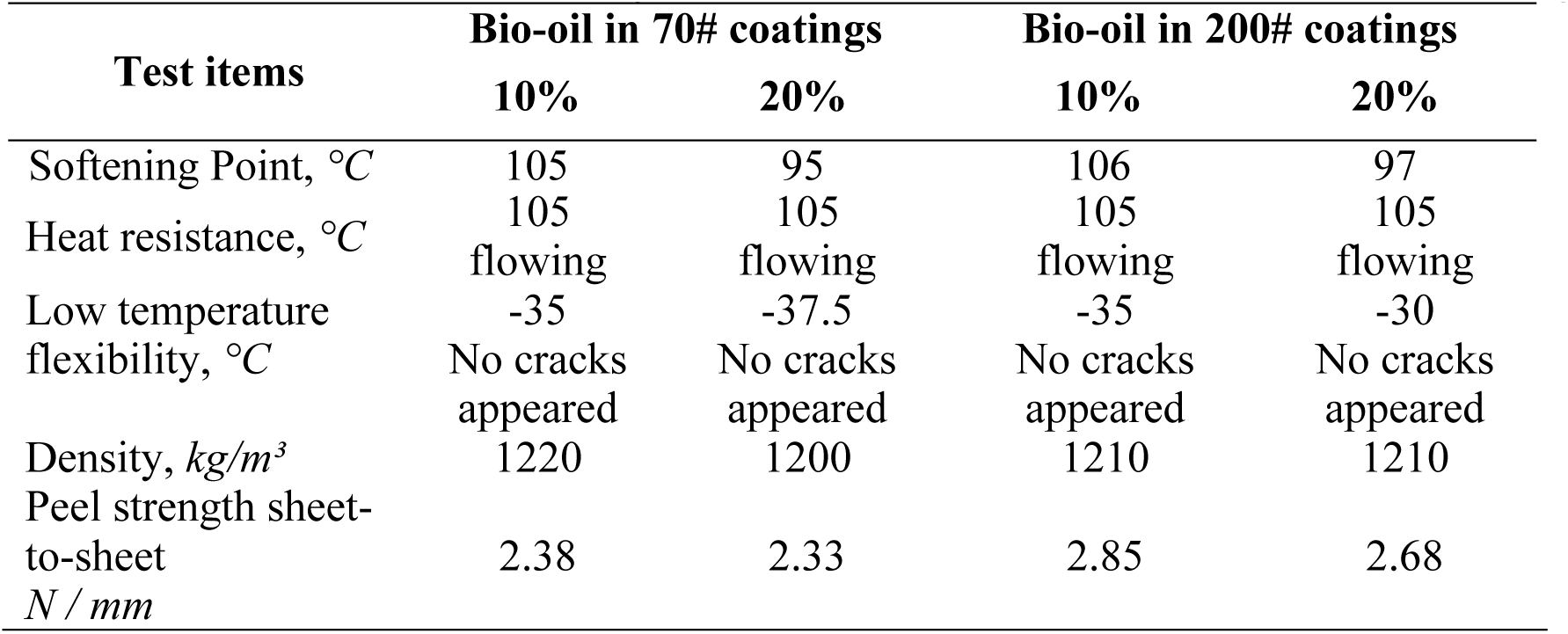
Effect of bio-oil added to grade 70# and 200# bitumen on SBS hot-melt coating sheet

In summary, the incorporation of pyrolysis bio-oil improves the softening point and peel strength of bio-bitumen, and its low-temperature flexibility is also improved when applied to self-adhesive sheet. In addition, it slows down the aging rate of bio-bitumen. Bio-oil exhibits a good alternative to coating sheet bitumen. Currently, the bio-oil is mainly investigated for road pavement asphalt, however, it still has great challenges, because the pavement asphalt requires high temperature stability and high mechanical properties due to the need to resist external loads, such as rutting resistance at high temperatures. However, the up-to-date research results show that the incorporation of most types of bio-oil adversely affects the rutting resistance at high temperatures (Yang et al. **2017**; Su et al. **2018**; Aziz et al. **2015**; Ingrassia et al. **2019**; Wen et al. **2013**). Although Wood based bio-oil could improve high temperature performance to some extent, it is not conducive to low temperature cracking performance (Yang et al. **2013**; Wen et al. **2013**). In comparison, bitumen coating sheet has high requirements on the viscosity and peel strength of bio-bitumen. Moreover, the coating sheet does not need to bear more external loads, which is undoubtedly more advantageous for the application of bio-bitumen. Therefore, the application of bio-oil in bitumen coating sheet shows better adaptability.

## CONCLUSIONS

The bio-oil produced by pyrolysis of wheat straw was applied to two different bitumen sheets. The effect of the bio-oil incorporation was studied by examining the physical properties. Results demonstrate that the addition of bio-oil increases the softening point and peel strength of the two coating sheets. Meanwhile, it is beneficial to the low temperature performance of self-adhesive coating. Interestingly, the small amount (about 8%) of water contained in bio-oil does not affect the performance of the bitumen sheet. This may be due to the presence of hydrogen bonds to help maintain structural stability. Except the substitution of 70# or 200# bitumen, the softening point of SBS hot-melt coatings decreased significantly with the increase of the substitution ratio. Compared to pavement bitumen, the application of bio-oil in coating sheet bitumen shows better adaptability. This study demonstrates the feasibility of using bio-oil in coating sheet materials. The results facilitate future applications of the method in coating sheet of buildings and roads.

## ACKNOWLEDGEMENTS

This study was funded by the National Natural Science Foundation of China (grant number 21676016, 21861132017, 21838001), and the State Key Laboratory of Special Functional Waterproof Materials (No. SKLW2018001).

## ABOUT THE AUTHOR

**Yanru Ren** is a graduate student at Beijing University of Chemical Technology, Beijing. China.

**Lei Zhang** is a polymer chemist at State Key Laboratory of Special Functional Waterproof Materials, Beijing Oriental Yuhong Waterproof Technology Co. Ltd, PR China.

**Wenfeng Duan** is the director of State Key Laboratory of Special Functional Waterproof Materials, Beijing Oriental Yuhong Waterproof Technology Co. Ltd, PR China.

**Zhongqiang Han** is a material engineer at State Key Laboratory of Special Functional Waterproof Materials, Beijing Oriental Yuhong Waterproof Technology Co. Ltd, PR China.

**Jia Guo** is a polymer chemist at State Key Laboratory of Special Functional Waterproof Materials, Beijing Oriental Yuhong Waterproof Technology Co. Ltd, PR China.

**Michael D. Heydenrych** is a chemical engineer at Department of Chemical Engineering, Faculty of Engineering, Built Environment and Information Technology, University of Pretoria, Republic of South Africa.

**Aijun Zhang** is an engineer National Energy Biorefinery Research and Development Center, Beijing University of Chemical Technology, Beijing, 100029, PR China.

**Kaili Nie** is a chemical engineer at National Energy Biorefinery Research and Development Center, Beijing University of Chemical Technology, Beijing, 100029, PR China.

**Tianwei Tan** is a chemical engineer at National Energy Biorefinery Research and Development Center, Beijing University of Chemical Technology, Beijing, 100029, PR China.

**Luo Liu** is a biochemical engineer at National Energy Biorefinery Research and Development Center, Beijing University of Chemical Technology, Beijing, 100029, PR China.

## REFERENCES

Akhtar, J., Kuang, S.K., and Amin, N.A.S. 2010. Liquefaction of empty palm fruit bunch (EPFB) in alkaline hot compressed water. Renewable Energy. 35:1220–1227. doi:10.1016/j.renene.2009.10.003

Aziz, M.M.A., Rahman, M.T., Hainin, M.R., and Bakar, W.A.W.A. 2015. An overview on alternative binders for flexible pavement. Constr. Build. Mater. 84:315–319. doi:10.1016/j.conbuildmat.2015.03.068.

Brennan, L., Owende, P. 2010. Biofuels from microalgae - A review of technologies for production, processing, and extractions of biofuels and co-products. Renewable Sustainable Energy Rev. 14:557–577. doi:10.1016/j.rser.2009.10.009.

Bridgwater, A.V. Peacocke, G.V.C. 2000. Fast pyrolysis processes for biomass. Renewable Sustainable Energy Rev. 4:1–73. doi:10.1016/S1364-1321(99)00007-6.

Capunitan, J.A., Capareda, S.C. 2013. Characterization and separation of corn stover bio-oil by fractional distillation. Fuel. 112:60–73. doi:10.1016/j.fule.2013.04.079.

Demirbas, A. 2001. Carbonization ranking of selected biomass for charcoal, liquid and gaseous products. Energy Convers. Manage. 42:1229–1238. doi:10.1016/S0196-8904(00)00110-2.

Fini, E.H., Kalberer, E.W., Shahbazi, A., and Basti, M. 2011. Chemical Characterization of Bio-binder from Swine Manure: Sustainable Modifier for Asphalt Binder. J. Mater. Civ. Eng. 23:1506–1513. doi:10.1061/(asce)mt.1943-5533.0000237.

Fernandez, A., Soria, J., Rodriguez, R., Baeyens, J., and Mazza, G. 2019. Macro-TGA steam-assisted gasification of lignocellulosic wastes. J. Environ. Manage. 233:626–635. doi:10.1016/j.jenvman.2018.12.087.

Goyal, H.B., Seal, D., and Saxena, R.C. 2008. Bio-fuels from thermochemical conversion of renewable resources: A review. Renewable Sustainable Energy Rev. 12: 504–517. doi:10.1016/j.rser.2006.07.014.

Ingrassia, L.P., Lu, X., Ferrotti, G., and Canestrari, F. 2019. Renewable materials in bituminous binders and mixtures: Speculative pretext or reliable opportunity? Resour., Conserv. Recycl. 144:209–222. doi:10.1016/j.resconrec.2019.01.034.

Kong, W., Baeyens, J., De Winter, K., Urrutia, A.R., Degrève, J., and Zhang H. 2019. An energy-friendly alternative in the large-scale production of soybean oil. J. Environ. Manage. 230:234–244. doi:10.1016/j.jenvman.2018.09.059.

Kong, W., Baeyens, J., Qin, P., Zhang H., and Tan, T. 2018. Towards an energy-friendly and cleaner solvent-extraction of vegetable oil. J. Environ. Manage. 217:196–206. doi:10.1016/j.jenvman.2018.03.061.

Kong, W., Miao, Q., Qin, P., Baeyens, J., and Tan, T. 2017. Environmental and economic assessment of vegetable oil production using membrane separation and vapor recompression. Front. Chem. Sci. Eng. 11:166–176. doi:10.1007/s11705-017-1616-4.

Önal, E., Uzun, B.B., and Pütün, E.A. 2011. Steam pyrolysis of an industrial waste for bio-oil production. Fuel Process. Technol. 92:879–885. doi:10.1016/j.fuproc.2010.12.006.

Prayogo, C., Jones, J.E., Baeyens, J., and Bending, G. 2014. Impact of biochar on mineralisation of C and N from soil and willow litter and its relationship with microbial community biomass and structure. Biol. Fertil. Soils. 50:695–702. doi:10.1007/s00374-013-0884-5.

Raouf, M.A., Williams, R.C. 2010. Rheology of fractionated cornstover bio-oil as a pavement material. Int J Pavements. 9:58–69.

Sanna, A., Li, S., Linforth, R., Smart, K.A., and Andresen, J.M. 2011. Bio-oil and bio-char from low temperature pyrolysis of spent grains using activated alumina. Bioresour. Technol. 102:10695–10703. doi:10.1016/j.biortech.2011.08.092.

Su, N., Xiao, F., Wang, J., Lin, C., and Amirkhanian, S. 2018. Productions and applications of bio-asphalts – A review. Constr. Build. Mater. 183:578–591. doi:10.1016/j.conbuildmat.2018.06.118.

Van de Velden, M., Baeyens, J., Brems, A., Janssens, B., and Dewil, R. 2010. Fundamentals, kinetics and endothermicity of the biomass pyrolysis reaction. Renewable Energy. 35:232–242. doi:10.1016/j.renene.2009.04.019.

Van de Velden, M., Baeyens, J., and Boukis, I. 2008. Modeling CFB biomass pyrolysis reactors. Biomass Bioenergy. 32, 128–139. doi:10.1016/j.biombioe.2007.08.001.

Van de Velden, M., Baeyens, J. 2006. A future “topper” in powder processing: Fast pyrolysis of biomass in a Circulating Fluidized Bed (CFB). Powder Handling and Processing. 18:354–360.

Wen, H.; Bhusal, S., Wen, B. 2013. Laboratory Evaluation of Waste Cooking Oil-Based Biobitumen as a Sustainable Binder for Hot Mix Asphalt. J. Mater. Civ. Eng. 25:1432–1437. doi:10.1061/(ASCE)MT.1943-5533.0000713.

Yang, X., You, Z., Dai, Q., and Mills-Beale, J. 2014. Mechanical performance of asphalt mixtures modified by bio-oils derived from waste wood resources. Constr. Build. Mater. 51:424–431. doi:10.1016/j.conbuildmat.2013.11.017.

Yang, X., You, Z.P., and Dai, Q.L. 2013. Performance Evaluation of Asphalt Binder Modified by Bio-oil Generated from Waste Wood Resources. Int. J. Pavement Res. Technol. 6:431–439. doi:10.6135/ijprt.org.tw/2013.6(4).431.

Yang, X., Mills-Beale, J., and You, Z. 2017. Chemical characterization and oxidative aging of bio-bitumen and its compatibility with petroleum asphalt. J. Cleaner Prod. 142:1837–1847. doi:10.1016/j.jclepro.2016.11.100.

Zargar, M., Ahmadinia, E., Asli, H., and Karim, M.R. 2012. Investigation of the possibility of using waste cooking oil as a rejuvenating agent for aged bitumen. J. Hazard. Mater. 233-234: 254–258. doi:10.1016/j.jhazmat.2012.06.021.

Zheng, J.L., Kong, Y.P. 2010. Spray combustion properties of fast pyrolysis bio-oil produced from rice husk. Energy Convers. Manage. 51:182–188. doi:10.1016/j.enconman.2009.09.010.

